# Human-like meaning maps from single-prompt VLM ratings of local scene meaning

**DOI:** 10.64898/2026.08.30.748100

**Authors:** Katarzyna Jurewicz, Yohaï-Eliel Berreby, B. Suresh Krishna

**Affiliations:** Department of Physiology, McGill University, Montreal, QC, Canada; Department of Philosophy, Centre for Cognitive Sciences, Jagiellonian University, Krakow, Poland; Mila—Quebec Artificial Intelligence Institute, Montreal, QC, Canada; UNIQUE Centre, Quebec Neuro-AI Research Centre, 3744 Jean-Brillant, Montreal, QC, Canada; Department of Ophthalmology, McGill University, Montreal, QC, Canada; Centre for Interdisciplinary Research in Music, Media and Technology, Montreal, QC, Canada

## Abstract

Visual attention is shaped both by low-level perceptual features and by the semantic properties of scene regions. However, compared to the salience of low-level image features, the characterization of image properties that lead to scene understanding has been more difficult. The semantic properties of scene regions have been operationalized via meaning maps, proposed by Henderson and Hayes (2017), by having human observers rate the informativeness of scene patches. However, this rating procedure is labor-intensive and limits the method’s versatility. An automated alternative, the DeepMeaning model, reduces rating costs but still requires human training data from a similar class of images, limiting generalizability. Here, we tested whether instruction-following vision-language models (VLMs) can generate human-like ratings in a zero-shot manner: using the original human-rating instructions as a single prompt without any task-specific training. We obtained meaningfulness ratings from both open-weight and proprietary VLMs for scene patches across three meaning-map datasets, and compared the resulting zero-shot AI meaning maps (AIMMs) to human meaning maps and maps obtained from the DeepMeaning model. Zero-shot AIMMs closely approximate human meaning maps, and maps from the task-specific DeepMeaning model, showing that VLMs can replicate human semantic judgments without any training. Zero-shot AIMMs demonstrate utility and validity, matching human meaning maps in their ability to predict human fixations and reproducing the same pattern of relative performance against other gaze-prediction models. We provide open-source code that allows researchers to use this zero-shot approach to quickly and flexibly explore meaning maps across spatial scales, prompts, and scene categories.

## Introduction

Understanding how scene salience and meaning shape our perception and visual behavior remains a central question in vision research. Early computational accounts emphasized low-level image features — local contrast, luminance, edge density — as primary drivers of fixation selection (Itti et al., 1998; Torralba et al., 2006; Wolfe, 2014). Other work demonstrated, however, that semantic content exerts a strong and largely independent influence on where observers look, with fixations clustering on regions that are meaningful, recognizable, or contextually informative (Ehinger et al., 2009; Nuthmann & Faul, 2026; Mack & Eckstein, 2011; Pedziwiatr et al., 2023; Yarbus, 1967). This dual influence of low-level salience and high-level meaning has stimulated the development of computational measures capable of capturing each in a spatially explicit form.

One influential approach for quantifying the spatial distribution of semantic content in scenes is the meaning maps method, introduced by Henderson and Hayes (2017). In this procedure, human participants rate isolated circular patches sampled from a scene on their perceived informativeness and recognizability — jointly referred to as “meaningfulness” — and these ratings are aggregated into a continuous spatial map. The approach offers an index of local semantic content that is analogous in form to low-level saliency maps, enabling direct comparison. While there is an ongoing discussion on the exact factors contributing to meaningfulness judgements (Leemans et al. 2024; Hayes and Henderson, 2022, 2025; Pędziwiatr et al. 2021, 2022), the practical relevance of meaning maps is supported by the observation that meaning maps can predict human fixation as well as, and sometimes better than, saliency models based on low-level image features; meaning maps can also explain a part of gazing behavior independent of the low-level salience (e.g. Hayes & Henderson, 2020; Henderson & Hayes, 2017, 2018; Peacock et al., 2019; Peacock, Hall, et al., 2023; Peacock, Singh, et al., 2023; for discussion see Nuthmann & Faul, 2026). The method has since been applied across a range of scene categories and contexts, establishing meaning maps as an important tool in the study of semantic contributions to visual attention (Bainbridge et al., 2019; Henderson et al., 2019; Oakes et al., 2024; Soyuhos et al., 2026).

Despite its utility, the human rating procedure underlying meaning maps and other subjective-rating maps is resource-intensive. Each image is decomposed into hundreds of patches, and each patch must be rated by multiple observers — a requirement that hinders this method’s applicability to very large image datasets and constrains the range of stimuli, questions, and spatial configurations that can be practically explored. Vision-Language Models (VLMs) — large neural networks trained on extensive image-text corpora — have recently emerged as a candidate class of models for probing human-like semantic representations. Hayes and Henderson (2025) proposed a partial solution for automatizing the construction of meaning maps in the form of DeepMeaning: a model that uses a pretrained VLM (Contrastive Captioner, CoCa; Yu et al., 2022) to extract features from scene patches, which are then used in a linear regression layer as predictors. The regression is trained with actual human ratings of the input patches to predict ratings for the new unseen ones. While DeepMeaning demonstrates strong performance, it requires human rating data from a similar image class for training. This limits its generalizability to different kinds of visual scenes and makes it inapplicable to new subjective-rating questions without additional surveying effort.

Instruction-following vision language models could provide a general-purpose alternative that avoids the need for such training data. Several lines of evidence suggest that VLMs develop internal representations and output patterns that align with human perceptual and cognitive judgments across diverse tasks (Marjieh et al., 2024; Nelson et al., 2025; Schlegel et al., 2025; Schulze Buschoff et al., 2025; Zhang et al., 2025), raising the possibility that meaningfulness ratings could be elicited from VLMs in a zero-shot manner — that is, through a natural-language prompt alone, without any task-specific fine-tuning or supervised training. Here, we investigated whether VLMs like the popular Google Gemini models can replicate human meaningfulness ratings of scene patches in a zero-shot procedure, i.e. without any additional model adjustments. We used images from three meaning map datasets: P21-scegram (Pedziwiatr et al., 2021), HH25-indoor and HH25-outdoor (Hayes & Henderson, 2025). To obtain a meaning map for an image we followed the meaning map procedure (Henderson and Hayes, 2017; Hayes & Henderson, 2025) and divided the image into patches. We submitted these patches to three models: the open-weight Gemma 4 31B IT and the proprietary models Gemini 2.5 Flash and Gemini 3 Flash Preview, along with the original instructions to humans in the meaning map procedure as the prompt. The resulting zero-shot AI meaning maps (AIMMs) were compared to human meaning maps (HMMs) and supervised DeepMeaning maps (DMMs), and tested for prediction of human fixations, along with other gaze-prediction models: (Graph-Based Visual Saliency model, GBVS (Harel et al., 2006); Adaptive Whitening Saliency model, AWS (Garcia-Diaz et al., 2012); and DeepGaze IIE, DGIIE (Linardos et al., 2021).

We demonstrate that meaning maps obtained zero-shot with VLMs closely approximate human meaning maps, establishing this approach as a practical tool for advancing research on the role of semantic content in visual attention. Its high throughput and high flexibility enables extensive exploration of spatial scale parameters, prompt formulations, scene categories, etc. Zero-shot AI subjective-rating maps make it practical to screen a large hypothesis space computationally. Beyond this methodological contribution, our findings carry an important implication for the broader meaning maps literature: given that AI-generated patch ratings bear striking similarity to human ones, researchers collecting meaning ratings through crowdsourced platforms can no longer assume that submitted responses originate from human observers, and appropriate safeguards should be considered in future data collection efforts.

## Methods

### Procedure for obtaining zero-shot AIMMs

The pipeline for generating zero-shot AIMMs followed the original one used for building human meaning maps (Hayes & Henderson, 2025; Henderson & Hayes, 2017), with the exception of the rating procedure, which was replaced by VLM-produced ratings. Each image was first segmented into partially overlapping, circular patches of two sizes: fine and coarse. Circular patches at the borders of the rectangular image extend outside the image. Patch diameter (in pixels) was different between the datasets (as detailed in the Datasets section below) and we kept these original patch sizes. The patches used by Hayes and Henderson (2025) were of the same pixel size as those used earlier (Henderson & Hayes, 2017). Patch sizes in P21-scegram (Pedziwiatr et al., 2021) were adjusted such that they matched sizes used in the original meaning map study (Henderson and Hayes, 2017) in degrees of visual angle (around 3 and 7 degrees of visual angle for fine and coarse patches respectively in both these studies) when viewed by human participants in the corresponding eye tracking experiments. To obtain meaningfulness ratings for all of the patches, we used three leading Vision-Language Models: Gemini 2.5 Flash, Gemma 4 31B IT, and Gemini 3 Flash Preview. Properties of these models and their performance on standard benchmarks (Yue et al., 2024, 2025; Rein et al. 2024) are summarized in **Supplementary Table S1**. We selected these models due to their performance-to-cost ratio and, in the case of Gemma 4 31B, open availability of model weights. Gemini 3 Flash Preview occupies the highest capability tier among the three; it falls short of the current frontier-class models such as Gemini 3.1 Pro, GPT-5.6 Sol, and Claude Fable 5, but its benchmark performance positions it as a strong option with a reasonable cost. Gemini 2.5 Flash, while remaining a performant and cost-efficient model, predates the Gemini 3 generation and is consequently surpassed on most benchmarks. Including Gemini 2.5 Flash allows for a partial assessment of whether advances in model capability between generations translate into measurable performance differences in “meaningfulness” ratings. Gemma 4 31B IT, released under an open-weight licence, represents one of the most capable freely deployable options and is competitive with mid-tier proprietary models.

Meaningfulness ratings were obtained by querying VLMs via the OpenRouter API (https://openrouter.ai/), which provides unified access to third-party vision-language models. The text of the instruction passed to the model was exactly the same as the one used for human subjects by Henderson and Hayes (2017) (the original instruction is available at https://osf.io/654uh/), except for the specification of the response format, which had to be adapted to text-based reporting. We did so by asking models to return a JSON-formatted “score” field, whereas human participants selected their answer on a visual Likert scale via mouse click. We asked the same question as originally posed: “*You will be presented with a patch of larger real-world scenes like the one above. Your task will be to rate how “meaningful” you think this scene patch is. What do we mean by “meaningful”? We want you to assess how “meaningful” an image is based on how informative or recognizable you think it is.*” (see the full text of the prompt in **Appendix**). Likewise, we used the sample examples of low- and high-meaning patches, which were embedded in the prompt as images. We asked for a single-value rating of “meaningfulness” on a 6-point scale. Each patch was rated independently. We set the model temperature parameter to 0, which provides maximally deterministic output.

A meaning map was then generated for each scene from the ratings, by setting each pixel’s value to the rating of the corresponding patch and averaging the rating data for pixels belonging to overlapping patches at each spatial scale separately and then averaging the two resulting maps together.

Importantly, it is highly unlikely that VLMs could obtain their ratings by having these values incorporated in the model from publicly available data. Even if the underlying photographs and meaning maps existed in training data, the models were never exposed to patch–rating pairs. VLMs evaluate patches independently, and reproducing a rating would require a complex, multi-step inference — matching the patch to its location in the original image, retrieving the corresponding meaning map, and reading off the rating at that location.

The full pipeline for running the zero-shot VLM rating procedure via the OpenRouter API is provided in an online repository (https://github.com/m2b3/aimm), together with step-by-step instructions.

### Datasets

To assess how well VLMs reproduce human meaning judgements, we used three open datasets of meaning maps and associated human fixation data.

#### P21-scegram

This dataset consists of patch ratings obtained online via Amazon Mechanical Turk (N = 69 raters, each patch rated by three unique raters), and human fixation data collected in the laboratory (N = 20 subjects) (Pedziwiatr et al., 2021, data available at https://zenodo.org/records/3490434). We converted raw ratings into meaning maps along the above-mentioned procedure and fixation data into fixation density maps using Gaussian smoothing (6 dB cut-off; Pedziwiatr et al., 2021). Stimuli used in this study come from the SCEGRAM database (Öhlschläger & Võ, 2017). For our purposes, we used only data from the “consistent” condition, in which scenes show objects that are typical for a given context (most suitable for the assessment of the local meaning; see Henderson et al., 2021; Pedziwiatr et al., 2021, 2022). This set consists of 36 photographs of real-world scenes. Each photograph was 688 x 524 pixels, and spanned ∼19.7 x 15 degrees of visual angle, when viewed by human subjects. When scenes were segmented into circular patches, fine patches (120 per image) had a radius of 53 px, coarse patches (48 per image) a radius of 123 px. Together, there were 168 patches per image.

#### HH25-indoor and HH25-outdoor

These datasets consist of a) human meaning maps; patch ratings were obtained from 1149 undergraduate university student raters with each patch rated by three unique raters) and b) human fixation density maps, from fixation data collected in the laboratory (N = 100 participants) (Hayes & Henderson, 2025; data available at https://osf.io/hcnfx/). Stimuli used for meaning map construction comprised the photographs used in previous studies on meaning maps (e.g. Henderson & Hayes, 2017; Henderson et al., 2018; Kiat et al., 2021) together with new photographs. Two images (‘target_greenhouse’ and ‘target_porch’) had different indoor/outdoor label across analyses in Hayes & Henderson (2025). Based on visual inspection, we (re-)assigned them to the indoor dataset, resulting in a total of 140 indoor (HH25-indoor) and 142 outdoor (HH25-outdoor) scenes. Subsets of those scenes (HH25-indoor, n = 51; HH25-outdoor, n = 49) were used by the original authors for testing fixation distributions. Each photograph was 1024 x 768 pixels, and spanned ∼35 x 25 degrees of visual angle when viewed by human subjects. When scenes were segmented into patches, fine patches (300 per image) had a radius of 43 px, and coarse patches (108 per image) a radius of 102 px. Together, there were 408 patches per image.

### Map comparisons

For comparing any two maps, following previous meaning map studies, we calculated Pearson’s linear correlation between the pixel-wise content of each map, resulting in one value of Pearson’s r per image. To summarize this distribution of r values across all images in a dataset, we applied the Fisher transformation to r values for individual scenes to obtain z values, calculated mean and 95% confidence interval (CI) from these z values, and transformed them back to r values. Means obtained this way were numerically very similar to median r values.

First, we used this procedure to compare zero-shot AIMMs to HMMs (**Figure 2)**, where we used raw, unnormalized and unsmoothed values of each map, as this is the most straightforward comparison for maps obtained with the same patches and same rating scale; our procedure included no histogram matching between the two maps, as in Hayes & Henderson (2025), but different from Henderson & Hayes (2017). We also used this procedure when comparing AIMMs to fixation density maps (**Figure 3A)**. To validate the use of AIMMs in the context of gazing behavior, we compared AIMM’s and HMM’s predictions of fixations to each other (**Figure 3B**), and to those from other gaze prediction models (**Figure 3A,C**): DeepMeaning (Hayes & Henderson, 2025), Graph-Based Visual Saliency model (GBVS; Harel et al., 2006), Adaptive Whitening Saliency model (AWS, Garcia-Diaz et al., 2012) and DeepGaze IIE (DGIIE; Linardos et al., 2021). The saliency maps from the AWS model were generated using the MATLAB code provided by the authors (http://persoal.citius.usc.es/xose.vidal/research/aws/AWSmodel.html). GBVS is distributed freely as MATLAB (https://github.com/Pinoshino/gbvs) and DGIIE (https://github.com/matthias-k/DeepGaze) as PyTorch (Paszke et al., 2019) implementations. For these analyses we smoothed the meaning maps (AIMM, HMM and DMM) to make them more comparable to other smooth maps (sigma = 15, as in Hayes & Henderson, 2025).

**Figure 1.**
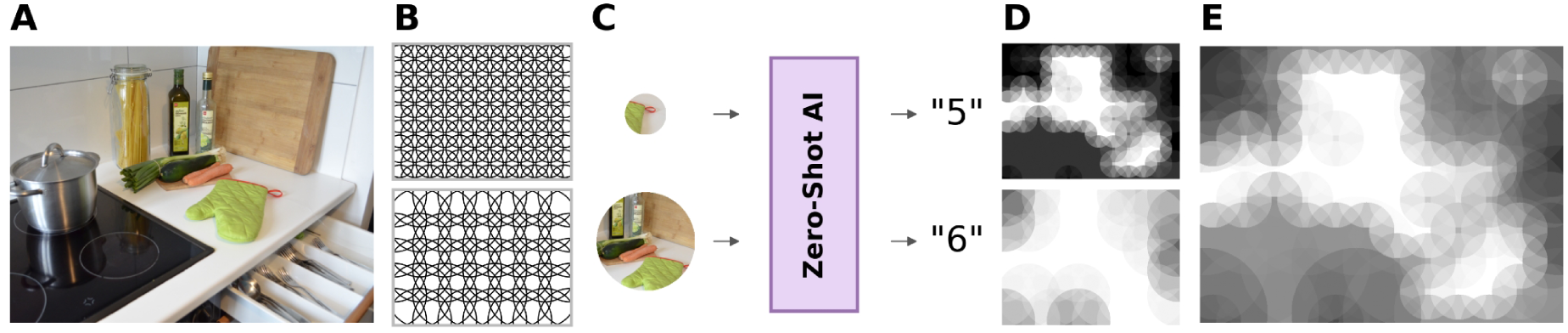
Illustration of the procedure for generating zero-shot AI meaning maps. **A)** The AI meaning maps were obtained for photographs of real-world scenes; example scene from P21-scegram dataset (Öhlschläger & Võ, 2017). **B)** A scene was decomposed into a series of overlapping circular patches at two spatial scales (fine and coarse). **C)** Each patch was passed to a VLM together with the prompt asking for meaningfulness rating on a scale of 1 to 6 (see the content of the prompt in **Appendix**). **D)** The ratings were transformed back into a 2D map separately for each scale (fine, coarse). **E)** The maps from two spatial scales were then averaged together.

**Figure 2.**
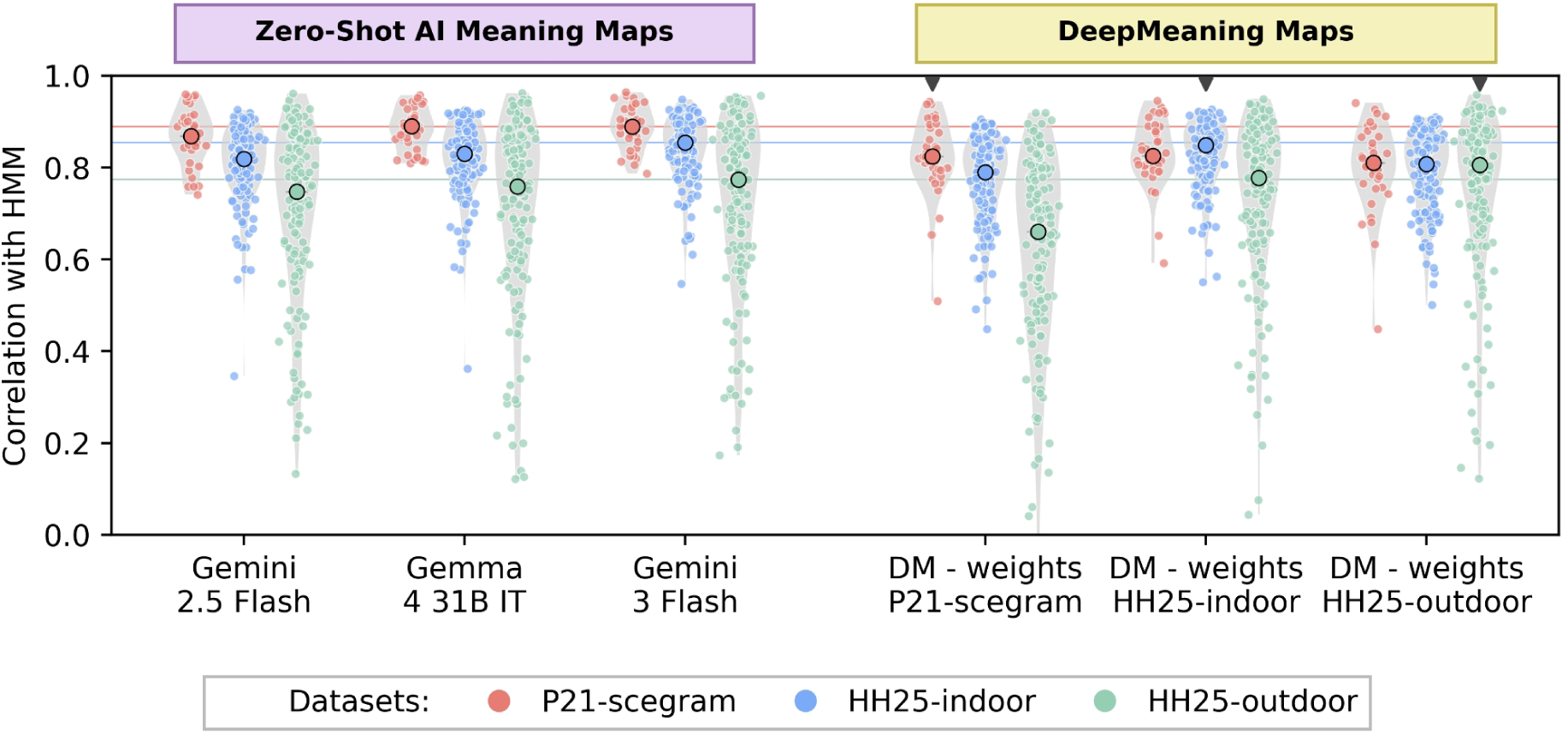
Recovery of human-like meaning maps from artificial neural networks. Meaning maps derived from zero-shot AI ratings are very similar to human meaning maps, and the level of similarity depends on the dataset. On the left we present correlation scores between human and zero-shot AI meaning maps for three selected models. Horizontal lines indicate average correlation obtained with the Gemini 3 Flash model. For comparison, on the right we present the results obtained with DeepMeaning’s linear-probing weights trained separately on each dataset, and evaluated on all three datasets. Small black triangles indicate same-set training and prediction, which used leave-one-scene-out cross-validation. Each dot represents one scene and larger, dark-circled dots indicate the median correlation score for the dataset (red: P21-scegram, blue: HH25-indoor, green: HH25-outdoor).

**Figure 3.**
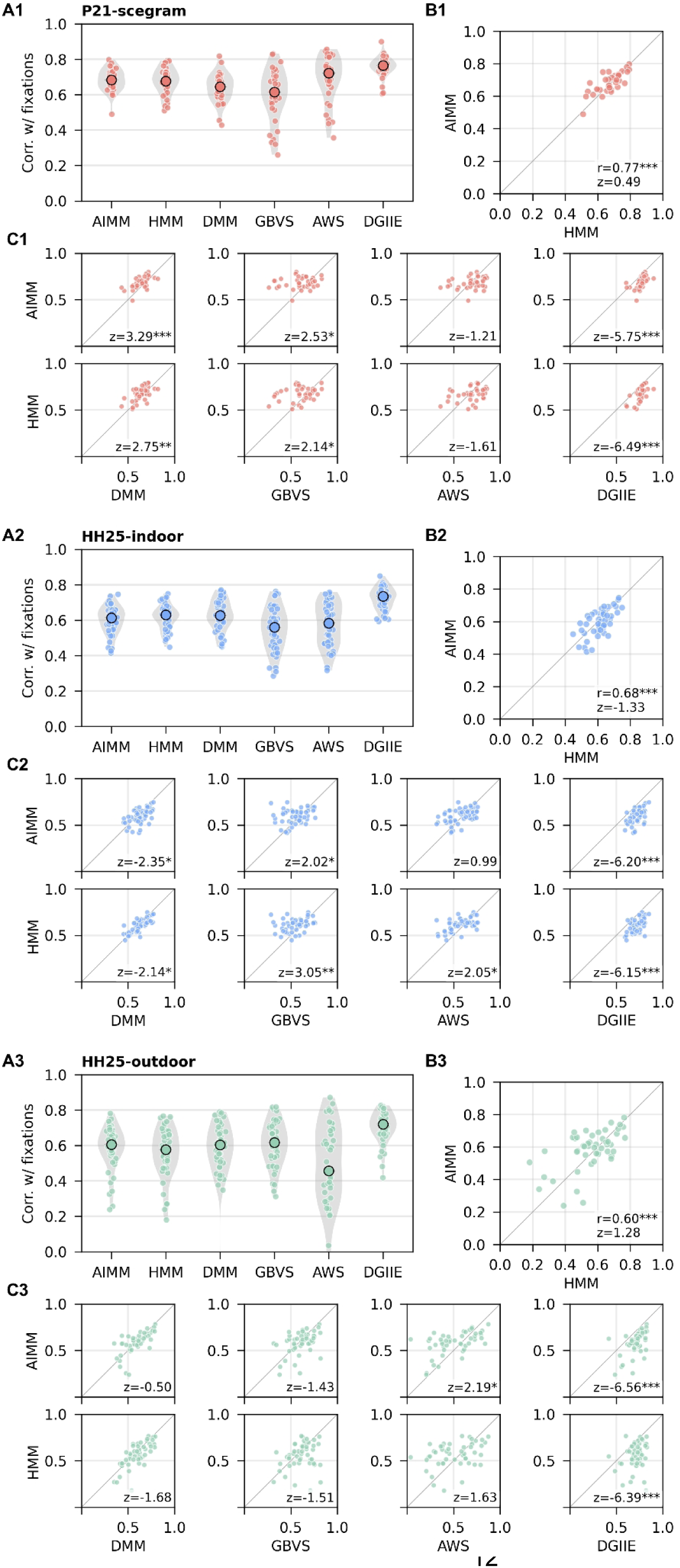
Zero-shot AI meaning maps predict distribution of human fixations similarly to human meaning maps. **A)** Correlations with human fixation density maps for meaning (AIMM, HMM, DMM), and gaze-prediction models (GBVS, AWS, DGIIE). Here, we consider zero-shot AIMMs that were obtained with Gemma 4 31B IT. DMMs were obtained with same-set training and prediction. Larger, dark-circled dots represent median values for each dataset, smaller dots represent single scenes. **B)** Direct comparison of pixel-wise correlations with fixations between AIMM and HMM shows high similarity of predictions. The r value represents a correlation coefficient between AIMMs’ and HMMs’ prediction of fixation density maps.The z value represents a mean difference between AIMMs’ and HMMs’ predictions. **C)** Similarly, both AIMMs and HMMs show corresponding patterns of predictions with other maps, pointing to the qualitative convergence between AIMMs and HMMs. Positive/negative z values indicate better/worse performance of AIMM and HMM compared to the other map. In B) and C) the reported correlation coefficients and z values were computed after Fisher z-transforming values on both axes. *** p < 0.001; ** p < 0.01; * p < 0.05.

Human fixation pattern demonstrates center bias, the tendency to fixate centrally. While this has no effect on comparisons between AIMMs, HMMs and DMMs, the presence or absence of bias has to be handled when we compare predictions of fixations. Unlike AIMMs, HMMs, DMMs and AWS, GBVS incorporates an explicit image-independent bias map, which implies that its salience estimates reflect a combination of image features and a general viewing tendency. Similarly, DGIIE by default uses a density bias map that was precomputed from MIT1003 human fixation density maps. To mitigate the role of image-independent bias in these maps for fair comparison, we ran GBVS with unCenterBias parameter = 1 and DGIIE with a uniform bias template of zeroes.

To compare whether any two maps, e.g. a AIMM and a HMM, correlated similarly on a pixel-wise basis with fixation density, we first z-transformed the Pearson’s r values from each map’s correlation with the fixation density map (e.g. AIMM with fixation density map and HMM with fixation density map, for each image). We then correlated these z values across images in each dataset, and again applied the Fisher z-transformation to the resulting r values. A mean and 95% CI was calculated on z values of this second-order correlation and these were then back-transformed to r values.

### Meaning maps from the DeepMeaning model

To situate our tests in the context of the currently available methods for generating meaning maps, we also applied the DeepMeaning approach proposed by Hayes and Henderson (2025) and compared its results with our zero-shot AIMMs. DeepMeaning model consists of two components: a pretrained Contrastive Captioner transformer (CoCa; Yu et al., 2022) that provides semantically rich latent features, and a linear model that is trained on a selected dataset to use these extracted features to predict human scene meaning ratings.

We generated the meaning maps with DeepMeaning code (Hayes & Henderson, 2025) for all datasets tested. Following the original report, for a model applied back to its own training dataset, each scene used the specific leave-one-scene-out cross-validation (LOSCV) fold that left that scene out. For a model applied to a different dataset, the prediction used the weights obtained by averaging coefficients and intercepts across all LOSCV folds on the source data (the ensemble weights). This is appropriate because none of the target scenes contributes to the source model’s training data.

Differences in numerical values reported here and in the original study are a consequence of using medians and z-transforming correlation coefficients for calculating means (rather than averaging raw r coefficients), performing correlations on unsmoothed data, and correcting the classification of one of the images. These changes were applied to align the processing of the results with other analyses, but did not change any qualitative patterns observed in the original report. For P21-scegram, DeepMeaning maps had not previously been computed.

Here, we needed to choose the size of the square grid for segmenting the scenes. If we matched the proportional area of a square patch (used by DeepMeaning) to the round patch (used for obtaining human ratings) used for HH25-indoor and HH25-outdoor, the square patch for P21-scegram would be 157 x 157 px. We assessed the role of the patch size on the outcomes with several patch sizes (88, 96, 128, 157 px), and, since the resulting differences were small and not systematic (**Supplementary Figure S1**), we used 128 px size for simplification and consistency with the patch size used in other datasets.

## Results

### Zero-shot AI meaning maps resemble human meaning maps

We first tested how well zero-shot VLM ratings could recover local scene meaning compared to human raters (**Figure 1**). We used several models to ensure that our results are not limited to the choice of one specific VLM. The models tested: Gemini 2.5 Flash, Gemma 4 31B IT, Gemini 3 Flash Preview all showed very high and dataset-dependent recovery of human-rated meaning maps, as verified at scene level (**Figure 2**; P21-scegram: r = 0.88, 0.89, 0.90, for the three models respectively; HH25-indoor: R = 0.82 / 0.83 / 0.85; HH25-outdoor: r = 0.75, 0.76, 0.77, see full report with 95% CI in **Table 1**).

**Table 1.** Correlation of human meaning maps and zero-shot AI meaning maps across models and datasets.

|  | P21-scegram | HH25-indoor | HH25-outdoor |
| --- | --- | --- | --- |
| Model | Mean [95% CI] ($r \rightarrow z \rightarrow r$) | | |
| Gemini 2.5 Flash | 0.88 [0.86, 0.90] | 0.82 [0.81, 0.83] | 0.75 [0.72, 0.78] |
| Gemma 4 31B IT | 0.89 [0.88, 0.91] | 0.83 [0.82, 0.84] | 0.76 [0.73, 0.79] |
| Gemini 3 Flash | 0.90 [0.88, 0.91] | 0.85 [0.84, 0.86] | 0.77 [0.74, 0.80] |
| DM - weights P21-scegram | 0.84 [0.81, 0.86] | 0.78 [0.77, 0.80] | 0.66 [0.62, 0.69] |
| DM - weights HH25-indoor | 0.85 [0.82, 0.87] | 0.84 [0.83, 0.85] | 0.77 [0.75, 0.80] |
| DM - weights HH25-outdoor | 0.82 [0.79, 0.85] | 0.80 [0.79, 0.81] | 0.79 [0.76, 0.81] |

The zero-shot VLM rating outcomes were systematically dataset- and model-dependent, with P21-scegram AIMMs having the highest, and HH25-outdoor AIMMs having the lowest similarity to HMMs across the 3 models (all p < 0.001, Mann-Whitney U, FDR-corrected). Differences between VLM models were smaller, but similarly consistent across the 3 datasets (with Gemini 3 Flash Preview obtaining the highest similarity to HMM and Gemini 2.5 obtaining the lowest similarity, from the models tested; all p < 0.001 for HH25-indoor and HH25-outdoor, all p < 0.06 for P21-scregram, Wilcoxon signed rank test, FDR-corrected). In addition, similarity of the ratings was more consistent in P21-scegram and HH25-indoor scenes than in HH25-outdoor scenes (HH25-indoor vs HH25-outdoor, all p < 0.001, P21-scegram vs HH25-outdoor, p ≤ 0.027; pairwise Fligner-Killeen test of variances, FDR-corrected) which is reflective of noisier human meaning ratings in outdoor scenes compared to indoor scenes and corroborates previous results (Henderson & Hayes, 2025). For reference, previously reported scene-level correlation observed between the two groups of subjects rating the same images was R = 0.87 (Hayes & Henderson, 2025). Thus, AIMMs performed at or close to the level of the noise ceiling of human raters.

We also compared the zero-shot VLM rating outcomes to an earlier use of VLMs in supervised map reconstruction, the DeepMeaning model (Hayes & Henderson, 2025). DeepMeaning is composed of two components: a pretrained Contrastive Captioner (CoCa) transformer (Yu et al., 2022) that is used as a feature extractor, and a linear model that is trained on a selected dataset to use these extracted features to predict human scene meaning ratings. When the linear model was trained on the same dataset (in a cross-validated procedure), its outcomes were similar to those that we obtained zero-shot (r = 0.84, 0.84. 0.79, for the three datasets respectively; **Figure 2**, results marked with black triangles; **Table 1** for the full report with 95% CI). DeepMeaning results for HH25-outdoor dataset were a bit better than in the best-performing zero-shot model (DeepMeaning vs Gemini 3 Flash Preview: HH25-outdoor, p = 0.040, Wilcoxon signed rank test, FDR-corrected), and worse than zero-shot for P21-scegram and r HH25-indoor (both p < 0.001). The better performance of DeepMeaning vs zero-shot VLM rating in HH25-outdoor data likely demonstrates the fitting-to-the-same-target-distribution advantage of DeepMeaning approach.

For the DeepMeaning model applied to a different dataset than the training one, all three sets of weights (obtained with P21-scegram, HH25-indoor and HH25-outdoor) produced lower results than the best-performing zero-shot model (p < 0.001), with the exception of HH25-indoor prediction with HH25-outdoor weights which were similar (p = 0.776).

This includes P21-scegram and HH25-indoor datasets cross-check, despite both datasets being composed of indoor scenes (both p < 0.001). This was not due to the specific choice of P21-scegram grid size (**Supplementary Figures S1 and S2**). The worse performance of DeepMeaning with P21-scegram weights on all datasets is likely because this dataset may be too small for effective training of the weights (n = 36 vs n = 140 and n = 142 in the other two datasets).

### Zero-shot AI maps and human meaning maps predict fixations equally well

Next, we evaluated how zero-shot AIMMs predict human fixations, compared to HMMs as well as other models used for predicting human fixations: DeepMeaning, as the closest counterpart, GBVS and AWS as the two models used for assessing the role of low-level saliency in guiding fixations, and DGIIE, state-of-the-art gaze-prediction model). This evaluation tests the ability of AIMMs to substitute for human-rated meaning maps in the context where meaning maps are primarily used, and an additional way for validating qualitative similarity of AIMMs and HMMs. We present these results using Gemma 4 31B IT. While its score was not the highest out of the three zero-shot models we tested, it is an open-weight model (i.e. its parameters are public), which facilitates future replicability of our results (results obtained with other models are qualitatively very similar and can be found in our online data repository).

Zero-shot AIMMs were strongly associated with where people looked in the scenes (**Figure 3A**; average correlation with fixation density maps, P21-scegram: R = 0.69; HH25-indoor: R = 0.61; HH25-outdoor: R = 0.60, see full report with 95% CI in **Table 2**). Importantly, this association was also very similar to the one of HMMs (**Figure 3B**; correlation between AIMM and HMM predictions of fixation density maps, P21-scegram: R = 0.77; HH25-indoor: R = 0.68; HH25-outdoor: R = 0.60, all p < 0.001; no significant difference between AIMM and HMM predictions of fixation density maps in any of the three datasets, p ≥ 0.183, Wilcoxon signed rank test). AIMMs and HMMs showed similar relationships to other types of fixation prediction (**Figure 3C**). Both AIMMs and HMMs predicted fixations better than DMMs within P21-scegram dataset (both p ≤ 0.006), worse than DMMs in H25-indoor dataset (both p ≤ 0.019), and not significantly different in H25-outdoor (both p ≥ 0.093). AIMMs also qualitatively replicated the results reported by Pedziwiatr et al. (2021), where human meaning maps predicted fixations worse than GBVS model (here, p < 0.012), but no different than AWS model (p = 0.226) and worse than DGIIE model (p < 0.001). While precise scores of different models varied between the datasets and comparisons (cf. Chakraborty & Zelinsky, 2022), the DGIIE model consistently outperformed other predictions.

**Table 2.** Correlation between meaning maps (HMM, AIMM and DMM), gaze-prediction models (GBVS, AWS, DGIIE) and human fixation density maps.

|  | AIMM | HMM | DMM<sup>1</sup> | GBVS | AWS | DGIIE |
| --- | --- | --- | --- | --- | --- | --- |
| Dataset | Mean [95% CI] (r→z→r) |  |  |  |  |  |
| **P21-scegram** | 0.69<br>[0.67, 0.71] | 0.68<br>[0.66, 0.70] | 0.65<br>[0.62, 0.67] | 0.62<br>[0.57, 0.66] | 0.71<br>[0.67, 0.75] | 0.76<br>[0.74, 0.78] |
| **HH25-indoor** | 0.61<br>[0.58, 0.63] | 0.62<br>[0.60, 0.64] | 0.64<br>[0.61, 0.66] | 0.56<br>[0.53, 0.60] | 0.59<br>[0.55, 0.62] | 0.73<br>[0.71, 0.75] |
| **HH25-outdoor** | 0.60<br>[0.56, 0.63] | 0.57<br>[0.53, 0.61] | 0.60<br>[0.55, 0.64] | 0.62<br>[0.58, 0.66] | 0.52<br>[0.45, 0.59] | 0.72<br>[0.69, 0.74] |

<sup>1</sup>DeepMeaning same-set training and prediction

## Discussion

In our study, we asked off-the-shelf vision-language models to produce zero-shot meaningfulness ratings of image patches, using the exact same instructions and examples as those originally given to human raters by Henderson and Hayes (2017). The resulting zero-shot AI meaning maps were highly consistent with human meaning maps, and predicted human fixations about as well as human maps do. These results were obtained using a natural-language and image-based prompt, without fine-tuning the underlying VLMs in any way, and without using a separately-trained decoder to interrogate the models’ latent space. Aside from the handful of examples provided in the description of the task to both humans and AI, our zero-shot methodology does not rely on any human-annotated data.

This last point is a critical difference between the zero-shot approach and the DeepMeaning model employed in the most closely related work (Hayes & Henderson, 2025). To predict human meaningfulness ratings DeepMeaning trains separate linear decoders for each dataset, with a leave-one-out procedure where the linear decoder is fit to all images except the one being predicted. Despite this relative advantage of being trained on images similar to the one being predicted, DeepMeaning does not consistently outperform zero-shot VLM ratings. Further, using DeepMeaning requires having some initial data collected on a similar sample of images. To use it with new photographs, it is not clear how the set of “similar” images that gives the best performance should be identified. Finally, DeepMeaning relies on explicit fitting to human data from the 768-dimensional latent representations output by a pretrained vision-language encoder. As a result, it inherently requires a non-negligible amount of data collection from human participants and is subject to overfitting concerns.

Any subjective attribute of image regions can in principle be used to construct an appropriate set of subjective-rating maps, following the example set by meaning maps (Henderson and Hayes, 2017), graspability maps (Rehrig et al., 2020), emotional maps (Pilarczyk et al., 2021) and the like. To date, such maps could be produced by specifying the task using natural language — optionally paired with a handful of examples — and asking human participants to rate image patches. Our results show that this approach can be straightforwardly extended from human to AI respondents. While the present work only considered the specific case of meaning maps, we see the production of other kinds of subjective-rating maps from AI responses as a promising avenue for future research. The approach is particularly well-suited to exploratory research that includes alternative construct definitions or novel scene categories, and a large-scale production of subjective-rating maps where collecting human ratings would be prohibitive. Zero-shot AI maps make it practical to screen a large hypothesis space computationally first, validating the selected outcomes with human data at the end, thus inverting the traditional workflow.

From perceptual domains, through affective judgements, up to generalized cognitive abilities, LLMs and VLMs have been shown to reproduce patterns of human judgements. They have recovered such canonical structures as the color wheel and the pitch spiral (Marjieh et al., 2024), matched or exceeded human accuracy at identifying facial emotions (Nelson et al., 2025), and solved standard emotional-intelligence tests and constructed new ones of comparable quality (Schlegel et al., 2025). Binz et al. (2025) showed that a then-frontier LLM, Llama 3.1 70B, could be fine-tuned on a very large corpus of human responses to act as a foundation model for human cognition, successfully predicting human behaviour in a wide variety of experiments. Finally, the narrowing gap between human and machine response profiles was documented by deploying Turing-test-like behavioral comparisons across language and vision paradigms (Zhang et al., 2025). These behavioral alignment findings are complemented by representational alignment research, which suggests that large-scale pretraining on human-generated or human-centric data causes models to develop internal representational structure that is fundamentally similar to human conceptual organization — providing the substrate for human-like semantic judgments without task-specific training. For example, triplet odd-one-out judgments collected from LLMs and VLMs (several million triplet judgements across a few thousand natural objects) showed that their multi-dimensional embeddings exhibit semantic clustering closely resembling human mental representations (Du et al., 2025; Hebart et al., 2020), strong alignment with neural activity patterns in brain regions associated with object recognition (EBA, PPA, RSC, FFA), and interpretability in terms of human-meaningful features. Thus, multidimensional representational geometry underlying human similarity judgments appeared to be recapitulated in models exposed to sufficient linguistic and multimodal data.

The meaning maps procedure asks people to rate patches on informativeness and recognizability, which are inherently semantic properties, not low-level image statistics. Indeed, the entire point of the method introduced by Henderson and Hayes (2017) is that it captures something that saliency models based on contrast, edges, and luminance *cannot*. This naturally leads to a question: is linguistic supervision specifically responsible for VLMs’ capacity to approximate human semantic ratings? We speculate that language plays several roles in the context of this study, as a source of semantic structure and world knowledge obtained by the VLMs, but perhaps more importantly, as a means for querying them. It is useful to disambiguate these roles: representational alignment between vision-only AI models and primate brains and behavior has already been extensively documented (e.g. Bakhtiari et al., 2021; Mehrer et al., 2021; Raugel et al., 2026; Schrimpf et al., 2018, 2020; Storr et al., 2021; Yamins et al., 2014; Yamins & DiCarlo, 2016; Zhuang et al., 2021). A recent landmark study, which trained DINOv3 foundation models on image data alone (Raugel et al., 2026) revealed that the models develop representations aligned with those from human fMRI and MEG, and that this alignment grows with model size, training amount, and the use of human-centric images during training. Thus, human-like semantic structure can emerge from visual experience alone. However, unlike the VLMs that we consider in our approach, neither a vision-only model such as DINOv3 nor a simple contrastive captioner like CoCa can be provided with a task description in words; this limitation in turn motivates the use of separately-trained probes like those used in DeepMeaning, whose limitations we described above. Thus, in the case of subjective-rating maps, language’s primary role may be that of an interface, allowing experimenters to describe a task in words and read back an answer as they would with human participants.

A potential concern for the zero-shot approach is reproducibility, as proprietary models accessed via API are subject to ongoing updates and eventual deprecation. However, up to low-level floating-point nondeterminism, reproducibility is attainable through locally deployable, open-weight models: pinning a specific revision of an open-weight VLM provides a stable reference comparable to the static CoCa encoder used in DeepMeaning. To summarize, it is useful to consider these two approaches as occupying complementary positions in the methodological space. DeepMeaning, once trained, is lightweight, and anchored to human norms on a defined reference dataset. It is an appropriate tool when the research question is well-specified, and the relevant dataset overlaps with the training distribution. Our zero-shot VLM rating approach, by contrast, offers immediate deployability without training, prompt-level flexibility, and access to the accumulated world knowledge of large multimodal systems. In the three relatively small datasets we tested, zero-shot VLM rating performs similarly to DeepMeaning and remains competitive by itself, but it is possible that there are other datasets or image types where DeepMeaning may retain an advantage. In such cases, it is conceivable that the two approaches can be synthesized: zero-shot ratings can be used to label synthetic datasets for training a DeepMeaning-style read-out, thereby combining the flexibility of zero-shot querying with the computational efficiency of a trained linear model operating on pre-trained CoCa features.

The present findings carry a concrete methodological implication for the practice of collecting meaning maps via online crowdsourcing platforms. Meaning map studies have historically relied on crowdsourced patch ratings — most commonly obtained via Amazon Mechanical Turk or similar platforms — as a scalable substitute for laboratory-based human judgments. The validity of that substitution rests on the assumption that submitted ratings originate from human participants. That assumption is now under challenge. There’s an ongoing debate if, or, to what extent large language model-based agents can pass as human participants in online surveys, calling into question the reliability of self-report research (Gordon et al., 2026; Van der Stigchel et al., 2026; Westwood, 2025; Zhang et al., 2025). Not all behavioral tasks are equally susceptible to AI substitution: tasks requiring perceptual-motor precision, such as those yielding reaction time distributions, are at present more difficult to imitate convincingly, whereas purely evaluative responses are not (Chetverikov, 2026). Meaning ratings as subjective evaluative judgments belong to the latter class. Consistent with this, the present findings show that AI-generated ratings reproduce human meaning maps closely enough that their human origin can no longer be presumed. This situation leads to two immediate conclusions. First, online collection from human participants should either be supplemented with robust human verification measures — beyond standard attention checks, which have been shown to be easily bypassed (Westwood, 2025) — or avoided in favor of laboratory-based collection where participant identity can be confirmed. Second, the present pipeline offers a practical alternative: if the goal is to obtain meaning maps for stimulus sets where human-machine correspondence is sufficiently high, querying AI can simply replace crowdsourced ratings. It may be more efficient to use zero-shot VLM ratings directly, provided that the specific VLM and prompt configuration have been validated against available human-annotated data of the type reported here. In this sense, the threat that VLMs pose to online behavioral research is also, to an extent, its own solution.

In summary, we demonstrate that VLMs’ output patterns align with human subjective judgments about meaningfulness of scene regions. Beyond reproducing human ratings, zero-shot AI meaning maps prove functionally equivalent to human meaning maps in predicting where people fixated, including reproducing the same pattern of relative performance against other gaze-prediction models. We provide an easy-to-use tool that can substitute for costly, dataset-specific human meaning-rating experiments without any meaningful loss of validity. Because this approach requires only a natural-language task description and a handful of examples, as in standard human data collection, it offers a practical way to screen large hypothesis spaces computationally before validating selected outcomes with human data.

## Open Practices Statement

Code for the pipeline, analyses and figures, with instructions for running it on new images, is available at https://github.com/m2b3/aimm; a two-image worked example runs without further downloads. The VLM ratings and all maps we computed will be archived at Zenodo (<u>10.5281/zenodo.22165217</u>). The scene images, fixation data and human meaning maps are distributed by their respective authors: P21-scegram stimuli via the SCEGRAM database (Öhlschläger & Võ, 2017) its fixation and rating data via https://zenodo.org/records/3490434 (Pedziwiatr et al., 2021), and HH25 data via https://osf.io/hcnfx/ (Hayes & Henderson, 2025). The repository documents the source paths and processing steps needed to rebuild them. None of the experiments was preregistered.

## Acknowledgements

We sincerely thank the authors of the data, models and code referenced above for making them available. We also thank Marek A. Pedziwiatr for all the helpful suggestions.

This research was funded by a grant from the NSERC Discovery Research program (RGPIN-2022-05399) and Supplement (DGECR-2022-00321), a CIRMMT Agile award, and a computing resources grant from Calcul Quebec and the Digital Research Alliance of Canada to BSK; IVADO Postdoctoral Research Funding (PostDoc-2021a-8859659558_2), travel awards from the Vision Health Research Network and UNIQUE, and a Bourse d’excellence UNIQUE to KJ; Vision Sciences Research Network (VSRN) PhD Recruitment Scholarship, Unifying Neuroscience and Artificial Intelligence - Québec (UNIQUE) MSc and PhD Excellence Fellowships, a Centre for Applied Mathematics in Bioscience and Medicine (CAMBAM) Fellowship, a McGill Department of Physiology Max and Jane Childress Entrance Fellowship, a McGill Faculty of Medicine and Health Sciences Internal Studentship, and a NSERC CGRS-D fellowship (CGRS D 613218-2026) to YEB.

