## Appendix for "Human-like meaning maps from single-prompt VLM ratings of local scene meaning"

**A.** Prompt sent to VLM for evaluation. Prompt contains interleaved text and images: image placeholders are in quotes and they are replaced with actual images.

The purpose of this study is to gain a better understanding of how people perceive real-world visual scenes like this:

"example_scene.png" - Real-world visual scene example

You will be presented with a patch of larger real-world scenes like the one above. Your task will be to rate how "meaningful" you think this scene patch is.

What do we mean by "meaningful"? We want you to assess how "meaningful" an image is based on how informative or recognizable you think it is.

For example, here are two scene patches taken from the example scene above that would be very low meaning:

"low_meaning_example1.png" - Edge of sidewalk

"low_meaning_example2.png" - Sky above house

Without the example scene it would be difficult to recognize what either of these image patches are.

And here are two example patches that would be very high meaning.

"high_meaning_example1.png" - Window and roof

"high_meaning_example2.png" - Back of car

Both of these patches contain information that is easily recognized even without the example scene.

You will be asked to rate how "meaningful" you think the remaining scene patch is using a 6-point scale.

A rating of 1 means you think the scene patch is very low meaning like the sky example.

A rating of 6 means you think the scene patch is very high meaning like the car example.

You must respond ONLY with valid JSON.

Use exactly this JSON format:

```
{
    "score": <number>,
}
```

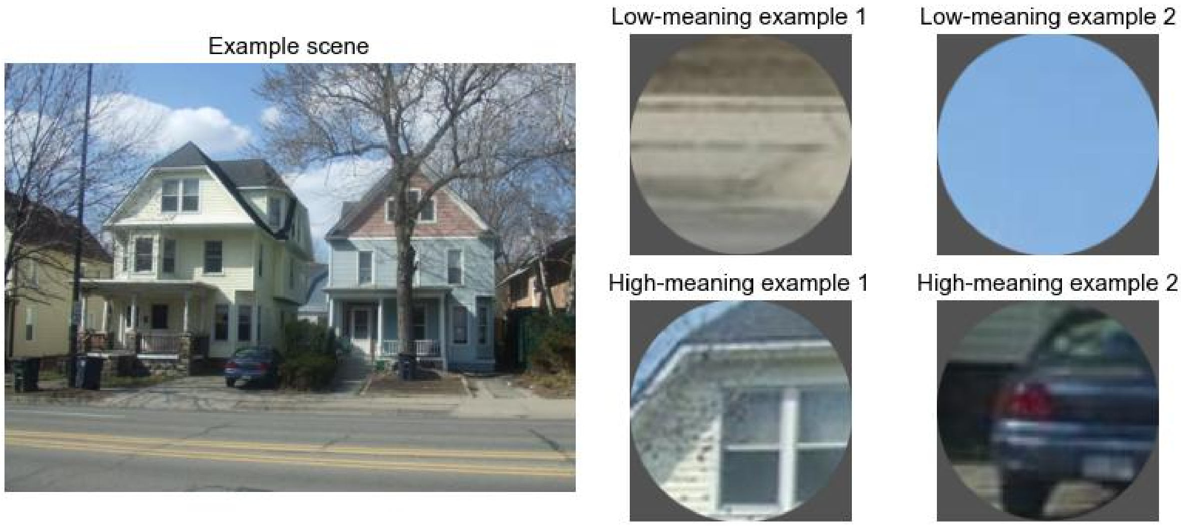
Example scene
Low-meaning example 1
Low-meaning example 2
High-meaning example 1
High-meaning example 2
