## Supplementary Materials for "Human-like meaning maps from single-prompt VLM ratings of local scene meaning"

### Supplementary Figures

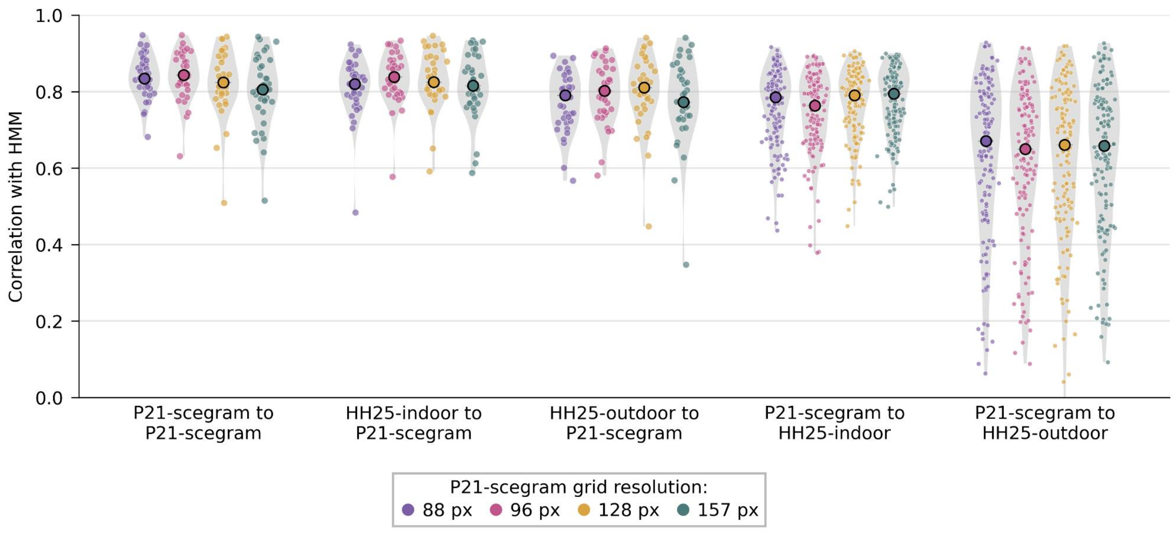


**Figure S1. Changing grid size has a limited influence on the results.** Relates to Figure 1. DeepMeaning results for P21-scegram are similar for different grid sizes applied to this dataset.

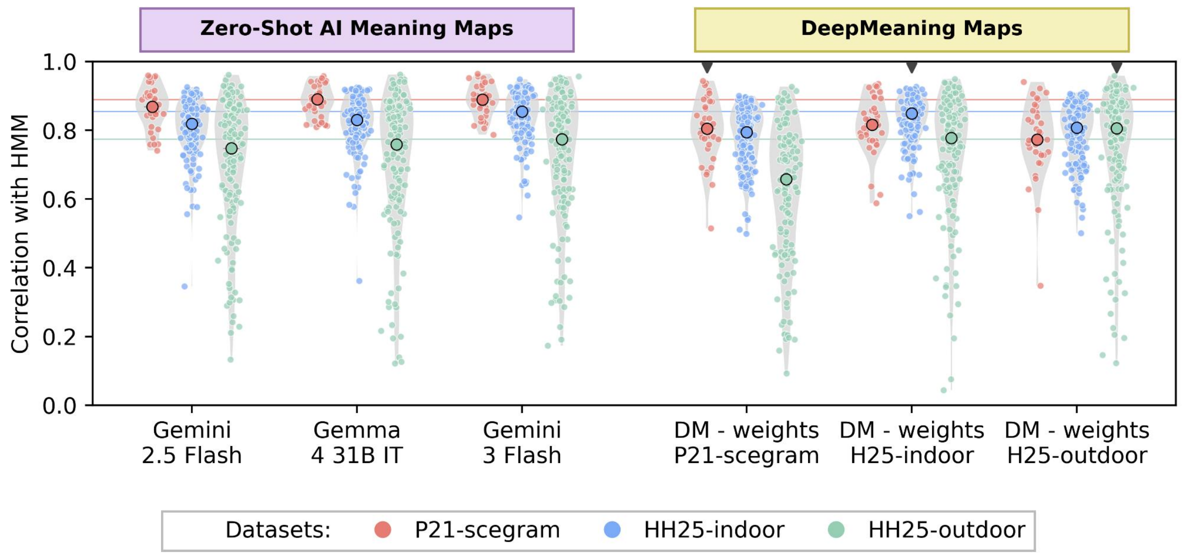


**Figure S2. Recovery of human-like meaning maps from artificial neural networks with DeepMeaning results using 157 px grid for P21-scegram.** Relates to Figure 1.

### Supplementary Tables

**Table S1**
Overview of AI models that were used in the study (Gemini 2.5 Flash, Gemma 4 31B IT, Gemini 3 Flash), and the frontier-class models as of writing (Gemini 3.1 Pro, GPT-5.6 Sol, Claude Fable 5), for reference.

| Model | Open weights? | Cost (in/out per 1M tokens)[#] | MMMU | MMMU-Pro | GPQA Diamond |
|---|---|---|---|---|---|
| **Gemini 2.5 Flash** | No | $0.30 / $2.50 | 79.7%[3] | n/a | 82.8%[3] |
| **Gemma 4 31B IT** | Yes | $0.13 / $0.38 | n/a | 76.9%[1, 3] | 84.3%[1] |
| **Gemini 3 Flash** | No | $0.50 / $3.00 | n/a | 87.6%[1] | 90.4%[1, 3] |
| **Gemini 3.1 Pro** | No | $2.00 / $12.00 | 77.5%[4] | 80.5%[1, 2, 3] | 94.3%[1, 2, 3] |
| **GPT-5.6 Sol** | No | $5.00 / $30.00 | 75.7%[4] | 83.0%[2, 3] | 94.6%[2] |
| **Claude Fable 5** | No | $10.00 / $50.00 | 74.3%[4] | 89.3%[4] | 94.1%[2] |

[#] Costs reported based on OpenRouter pricing as of writing.
[1] Google (DeepMind/Gemma technical reports & model cards)
[2] OpenAI — openai.com/index/gpt-5-6
[3] llm-stats.com — /benchmarks/mmmu, /benchmarks/mmmu-pro, /benchmarks/gpqa
[4] Vals AI, via benchmarklist.com
